# Influenza Classification from Short Reads with VAPOR Facilitates Robust Mapping Pipelines and Zoonotic Strain Detection for Routine Surveillance Applications

**DOI:** 10.1101/597062

**Authors:** J. A. Southgate, M. J. Bull, C. M. Brown, J. Watkins, S. Corden, B. Southgate, C. Moore, T. R. Connor

**Affiliations:** Organisms and Environment Division, School of Biosciences, Cardiff University, United Kingdom; Public Health Wales, University Hospital of Wales, Cardiff, United Kingdom; MRC Centre for Regenerative Medicine, University of Edinburgh, United Kingdom

**Keywords:** Influenza, Whole-genome sequencing, Classification, Bioinformatics, Surveillance

## Abstract

**Background:** Influenza viruses are associated with a significant global public health burden. The segmented RNA genome of influenza changes continually due to mutation, and the accumulation of these changes within the antigenic recognition sites of haemagglutinin (HA) and neuraminidase (NA) in turn leads to annual epidemics. Influenza A is also zoonotic, allowing for exchange of segments between human and non-human viruses, resulting in new strains with pandemic potential. These processes necessitate a global surveillance system for influenza monitoring. To this end, whole-genome sequencing (WGS) has begun to emerge as a useful tool. However, due to the diversity and mutability of the influenza genome, and noise in short-read data, bioinformatics processing can present challenges.

**Results:** Conventional mapping approaches can be insufficient when a sub-optimal reference strain is chosen. For short-read datasets simulated from influenza H1N1 HA sequences, read recovery after single-reference mapping was routinely as low as 90% for human-origin influenza sequences, and often lower than 10% for those from avian hosts. To this end, we developed a *de* Bruijn Graph (DBG)-based classifier of influenza WGS datasets: VAPOR. In real data benchmarking using 257 WGS read sets with corresponding *de novo* assemblies, VAPOR provided classifications for all samples with a mean of >99.8% identity to assembled contigs. This resulted in an increase in the number of mapped reads by 6.8% on average, up to a maximum of 13.3%. Additionally, using simulations, we demonstrate that classification from reads may be applied to detection of reassorted strains.

**Conclusions:** VAPOR has potential to simplify bioinformatics pipelines for surveillance, providing a novel method for detection of influenza strains of human and non-human origin directly from reads, minimization of potential data loss and bias associated with conventional mapping, and allowing visualization of alignments that would otherwise require slow *de novo* assembly. Whilst with expertise and time these pitfalls can largely be avoided, with pre-classification they are remedied in a single step. Furthermore, our algorithm could be adapted in future to surveillance of other RNA viruses. VAPOR is available at https://github.com/connor-lab/vapor.

## Introduction

Influenza viruses are enveloped, single-stranded, segmented negative-sense RNA viruses of the family *Orthomyxoviridae*. Influenza A and B have 8 genome segments encoding major structural and non-structural proteins, with major antigenic recognition sites within the two spike proteins haemagglutinin (HA) and neuraminidase (NA). Influenza replication in the host cell occurs by means of a viral encoded polymerase that lacks proof-reading capability, leading to frequent point mutations. Accumulation of these point mutations within the antigenic recognition sites of HA and NA can result in host immune evasion, thereby causing annual seasonal epidemics[1] [2]. Current estimates suggest that seasonal influenza A and B cause 4-5 million severe infections [3] in humans with approximately 291,000 to 645,000 [4] deaths per year globally.

Whilst influenza B remains largely a human pathogen, influenza type A is a zoonotic virus infecting a wide range of avian and other non-human species. To date 18 haemagglutinin and 11 neuraminidase types have been recognised, with a reservoir for the majority within birds[5]. These viruses have the capability to reassort leading to the emergence of new strains [6]. Should these include new HA and NA proteins, pandemic viruses can emerge that completely evade the host response leading to global epidemics of often high morbidity and mortality in both non-human and human hosts.

Influenza viruses therefore represent pathogens of major importance. In response, global laboratory and epidemiology networks have been established and managed by the World health Organisation (WHO) to monitor this constantly evolving viral landscape. The aim of these networks is to provide an early warning system for the emergence of seasonal viruses that respond less well to vaccination and to detect viruses with pandemic potential.

Whole genome sequencing (WGS) has been used to study the influenza virus genome for over a decade, and is emerging as an important tool in research and surveillance[7][8][9][10]. Protocols have been developed [11][12] that facilitate routine monitoring of isolates by public health organizations, as well as the study of transmission events [10][13]. Two important data sharing resources exist to this end; the NCBI Influenza Virus Resource (NIVR) [14], and the Global Initiative on Sharing All Influenza Data (GISAID) [15], wherein over a hundred thousand influenza genome segment sequences can be found at the time of writing, from isolates sequenced across the globe. Whilst methodologies exist involving passage of isolates, sequencing can be performed directly from clinical swabs with single-reaction genomic reverse transcription polymerase chain reaction (RT-PCR) [16][12]. Furthermore, bioinformatics pipelines have begun to be developed for efficient processing of this data [17][18].

Despite the increasing application of Next-Generation Sequencing (NGS) to influenza, the pitfalls associated with current mapping approaches have not been explored in depth. Influenza virus *de novo* assembly also poses additional challenges due to biological population complexity and additional error resulting from RT-PCR[19][20]. Firstly, we aim to provide evidence that current mapping approaches can, due to diversity of influenza genome sequences, routinely result in a large number of unmapped reads. In turn, this can potentially result in data loss and bias in sequences that are subsequently recovered, analyzed, and submitted to public databases. This has been previously noted in study of human immunodeficiency virus (HIV) [21]. Whilst alternatives, such as read classification by mapping to a large database of influenza sequences [22] and subsequent *de novo* assembly can help to resolve this issue, such pipelines are often complex, slow, and require expertise that is not necessarily available in routine surveillance. Secondly, even if bioinformatics pipelines are chosen judiciously, sequences of zoonotic origin may fail to be identified, resulting in a dataset that appears to be low coverage, missing segments, or missing potential future pandemic reassortments. Furthermore, even with recent assembly programs, misassembly can occur [21].

We aim to show that this problem can be resolved by classification of isolates from reads prior to analysis by directly querying a De Bruijn graph (DBG) built directly from Illumina sequencing reads. Mapping reads directly to a DBG has been previously argued to be less biased than that of mapping to assembled contigs [23]. Directly querying DBGs instead of assembled sequences has been previously addressed [23][24][25][26], although most previous work has focused on mapping reads to a DBG, and not diverse RNA virus sequence data. To our knowledge these approaches have not been applied to pathogen classification from reads. Instead of mapping reads to a DBG, we sought to further develop a simple method for querying short influenza genome sequences against a short read DBG in order to retrieve the most similar reference for mapping applications. In doing so, we leverage the large number of publicly available influenza segment sequences. We compare our tool that implements this algorithm, VAPOR, with both a slow BLAST-based [27] approach and fast kmer-based MASH [28], and show superior or equivalent results in several use cases with reasonable runtimes. We show through simulation, that given a set of influenza reads, possibly contaminated with human or bacterial sequences, a highly similar strain in the NIVR database (>20,000 strains) can be selected, achieving reasonably fast, and often near-strain-level, classification.

## Methodology

### WGS Datasets

Total RNA was extracted from patient samples using the NucliSens easyMAG instrument according to the manufacturer’s instructions. Following RNA extraction, a one-step RT-PCR (Quanta biosciences qScript XLT kit, following manufacturer’s instructions) was then undertaken to generate DNA for sequencing using the primers previously described for influenza A [12] and influenza B [11]. Sequencing was performed using Illumina sequencing instruments. Libraries were prepared using NexteraXT, and samples were then multiplexed for sequencing. Samples were run on a MiSeq (2×250bp V2 kit 44 samples) and NextSeq (2×150bp Medium Output kit 213 samples). In total, 257 samples were utilized. Short read data can be found at https://s3.climb.ac.uk/vapor-benchmark-data/vapor_benchmarking_realdata_reads_filtered_18_03_18.tar. For publicly available data, any reads that were classified as human by Kraken2 [29], or those that mapped to the hg38 human genome with minimap2[30], were removed.

These WGS datasets were then processed by extraction of influenza reads by mapping with minimap2 [30] to 8 curated influenza segment reference fasta files (19594 sequences in total), one at a time, produced by from all influenza segment sequences downloaded from the NIVR (https://www.ncbi.nlm.nih.gov/genomes/FLU/) and clustered to 99.5% identity with cd-hit-est [31]. Extracted reads were assembled with IVA [19]. For all 257 datasets used, a near-full length (>90%) contig could be assembled for at least one major segment protein. Samples for which a contig could not be assembled were not used. In total, 1495 segment contigs were included.

### Mapping Assessment

Four mapping programs were assessed in this analysis: Minimap2 [30], BWA-MEM [32], NGM [33], and Hisat2 [34]. Default settings were used for all tools. Each experiment can be reproduced using the code and instructions found at https://github.com/connor-lab/vapor_mapping_benchmarking. Four mapping simulations were performed in total.

For assessment of the sufficiency of single reference strains for mapping diverse samples, two simulations were performed. For assessment of robustness to species origin, read sets were simulated with Artificial-FastqGenerator (AFG)[35] from 552 avian, 16,679 human, and 4054 swine H1N1 HA coding sequences from the NIVR [14]. An additional 0.05% *in silico* substitution was introduced into simulated reads to account for RT-PCR technical errors and biological intrahost variation. This rate was chosen to be in accordance with experimental observations made by Orton *et al.* (2015) [20], although it may be conservative. Reads were then mapped to the A/California/07/2009 (H1N1) HA reference sequence. For assessment of robustness to divergence, technical and biological noise, reads were simulated from A/Perth/16/09 (H3N2) HA, with additional *in silico* mutation with per-base rates between 2% and 16%, which was performed uniformly across the chosen reference sequence; reads were simulated as above, then mapped back to A/Perth/16/09 (H3N2). This was performed 1000 times for each mutation rate. A/California/07/2009 (H1N1) and A/Perth/16/2009 (H3N2) were used as references since they are common clade representatives, as well as vaccine recommendations. Samtools [36] was used to retrieve successfully mapped reads, which were then counted.

For comparison of mapping with and without VAPOR classification, and potential zoonotic virus detection, 33,133 unique full-length influenza A HA coding sequences of any lineage or species were downloaded from the NIVR, and 5000 pairs were chosen randomly; the first of the pair was used for read simulation as above, and the second as a mapping reference. In the second run with VAPOR classification, a single sequence was randomly chosen as before, but the reference was chosen by VAPOR version 1.0.1. As before, successfully mapped reads were extracted with samtools, then counted.

To assess the potential benefit of classification with VAPOR on real data, 206 of 257 read pairs were subjected to mapping with Minimap2 with default settings for short reads (-x sr), both with and without VAPOR classification. 51 of 257 samples with less than 1000 HA reads were excluded to avoid very low coverage samples skewing calculation of mean percentage gain. In the first case, reads were mapped to a set of 4 HA references from different subtypes: A/Perth/16/2009 (H3N2), A/California/07/2009 (H1N1), B/Florida/4/2006 (Yamagata), B/Brisbane/60/2008 (Victoria). In the second case, VAPOR was used to choose a single reference from 53,758 influenza A and B HA references. The number of reads mapping and the number passing VAPOR pre-filtering was recorded in each case.

### Algorithm Overview

Figure 1 gives a simplified overview of the seed-and-extend algorithm used by VAPOR. As input, VAPOR takes a fasta file of full-length reference segment sequences, and a fastq (or fastq.gz) file of influenza WGS reads. Firstly, VAPOR builds a set of kmers *R* from the reference sequences. Next, for *i* ∈ *I*, where *I* is the number of reads, the *i*th read is decomposed into a set of non-overlapping words *A*_*i*_, and if |*A*_*i*_ *∩ R*| *\* |*A*_*i*_| *≤ t*, for some specified parameter *t*, the read is discarded. This is repeated for the reverse complement; if both are kept, the highest score decides orientation. Next, a weighted DBG (wDBG), 𝒲 = (*N, E, W*), where *N, E, W* are sets of nodes, edges, and weights, is built from the surviving reads; explicitly, edges are represented as overlapping kmers, and weights are kmer coverage in the remaining reads. Let *j* ∈ *J* be the index of the *j*th kmer. Any kmer *e*_*j*_ ∈ *E* with corresponding weight *w*_*j*_ ∈ *W* less than a coverage parameter *c* is discarded. Next, querying is performed. Let *m* ∈ *M* be the index of the *m*th reference sequence. Each reference sequence, *s*_*m*_, with length *L*_*m*_, is decomposed into a sequence of kmers. Querying proceeds in four phases, where the query is walked along the wDBG: kmer seeding, trimming, bridging, and scoring. Firstly, an array *a*_*m*_ is created from exact kmer matches, where *a*_*mn*_ is the weight of the *n*th kmer of the *m*th reference, and any kmers not in *E* are set to zero. In trimming, any gaps (sequence of zeros) in this array, representing kmers not present in the wDBG, is then expanded (seed trimming) to suboptimal branch points in the graph within *ρ* positions of the gap, in order to heuristically prevent suboptimal seeds to low coverage regions of the wDBG, possibly generated by error. Let *k* ∈ *K* be the index of the *k*th gap in *a*_*m*_. For any gap *g*_*k*_ of length *l*, a bridge *b*_*k*_ is formed by walking *l* locally optimal (where there is a branch, the edge with the highest weight) edges in the wDBG from the last matching base. Each bridge, *b*_*k*_, a string, is then compared to the *k*th gap string, the original substring in the reference sequence corresponding to the gap. For any matching base between the gap string and the bridge, the corresponding weight of the kmer with this base as its first position is inserted into *a*_*m*_ at the corresponding position. Finally, since the scores in *a*_*m*_ correspond to kmer weights, per-base scores are re-calculated. The score of any base (including the final *k -* 1 that do not have corresponding kmers) is given the highest score of all kmers that contain it in the sequence, excluding mismatching bridge positions. This is calculated using a deque. Finally, the total score is given by 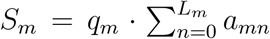 where *q*_*m*_ is the proportion of bases of *s*_*m*_ found in the graph. For speed considerations, only a subset of seed arrays are extended: those with a fraction of nonzero elements greater than a user-defined parameter `--min kmer cov` (default: 0.1), and in a top user-defined percentile `--top seed frac` (default: 0.2). VAPOR is implemented in Python3, with source code available at https://github.com/connor-lab/vapor.

**Figure 1:**
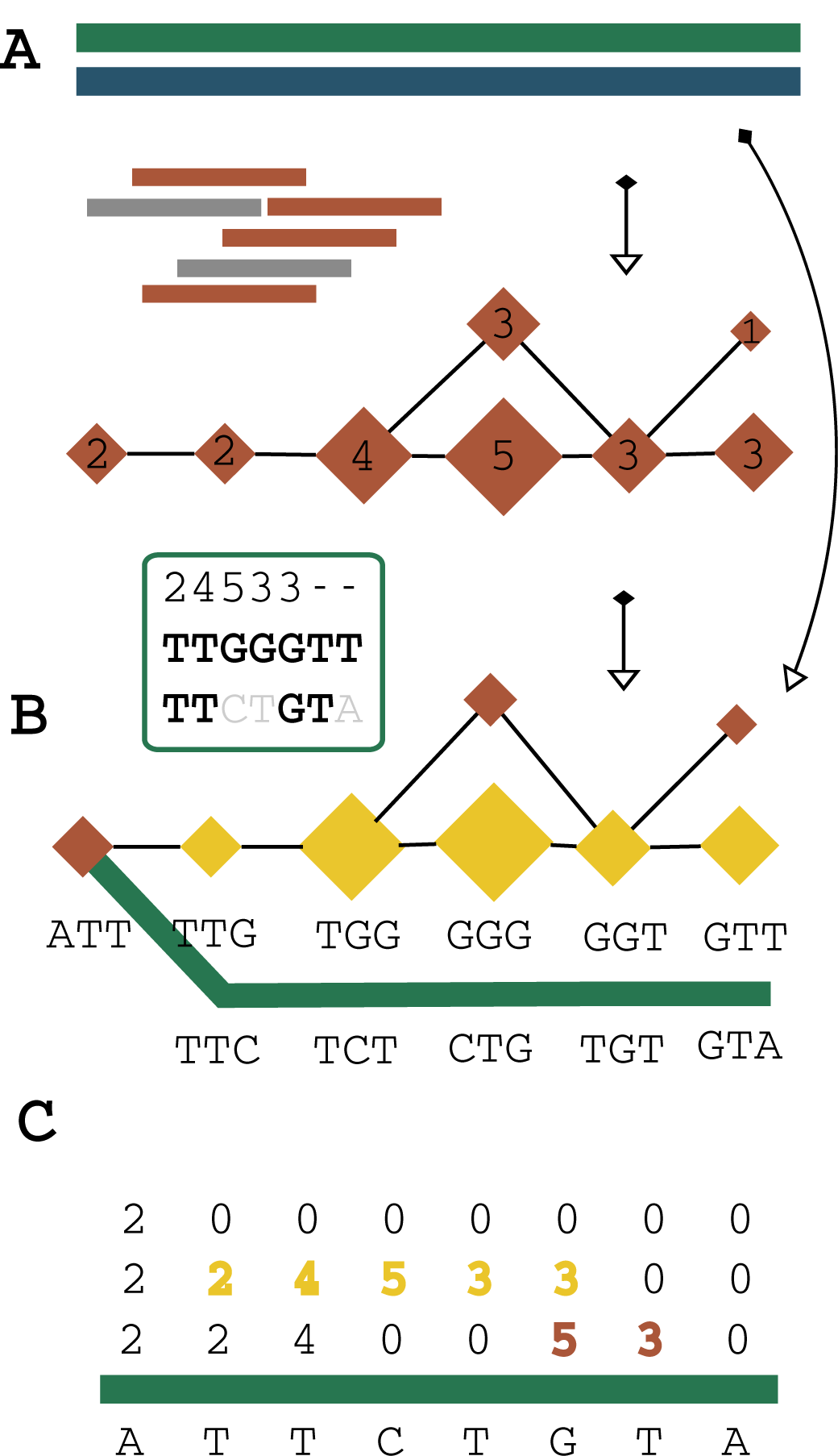
Simplified VAPOR algorithm overview. Firstly (A), contaminant reads (grey) are filtered out according to the fraction of shared kmers with the reference sequences (green, blue), and a wDBG is built from the surviving influenza reads (red). Sequences are queried against the wDBG in two main phases (B): kmer seeding and bridge extension. Exact kmer matches are used as seeds (ATT in the example). For a given gap of length *l* in the query kmer sequence (here *l* = 5), we attempt to traverse the graph *l* locally optimal steps in the wDBG to produce a bridge sequence (yellow) and corresponding scores. Comparison of the bridge and gap sequence is shown in B, with mismatching bases greyed out. When scoring (C), bridge scores are used to fill in gaps (yellow), mismatched bases are not counted, and the score of the kmer with the greatest weight that may cover a given base is used (red). Final arrays are summed and multiplied by the fraction of bases with nonzero coverage.

### Classification Benchmarking

VAPOR was compared to MASH [28] and BLAST [27] read classification by simulation. BLAST consensus classification was performed by BLASTing each read, taking the best scoring references by e-value then bit score, summing the number of times each result occurs in all reads, and returning the most frequent. Reads were simulated as follows: a reference, *s*_*o*_, was chosen from 46,724 unique full-length influenza A HA sequences from the NIVR, and mutated uniformly with a given probability (0.01, 0.02, 0.03) to generate a mutated sequence *s*_*m*_; reads were simulated with AFG as before, with a higher uniform error rate of 1%, in order to provide a challenging classification task representative of difficult datasets. To provide an additional challenge, we simulated an intra-host population with 4 minor sequences, mixed in the ratio of 100:5:1:1:1, with each minor sequence additionally mutated by 1% relative to the major sequence. This process was performed 500 times for each category. Performance was assessed as follows: The Levenshtein distance of the mutated sequence *s*_*m*_ was taken with respect to the original sequence *s*_*o*_ as a baseline, denoted by *L*(*s*_*m*_, *s*_*o*_); the reads were classified by each tool with all 32,804 references as a database, and the best hit *s*_*c*_ returned by each were compared to the mutated sequence to obtain *L*(*s*_*m*_, *s*_*c*_). Global alignment was performed with the pairwise2 module of Biopython[37] (with cost parameters 0, −1, −1, −1). We defined the additional Levenshtein distance, *L*_*A*_ = *L*(*s*_*m*_, *s*_*c*_) *- L*(*s*_*m*_, *s*_*o*_). This distance was chosen because, for mutated sequences, it captures the additional error in classification beyond that caused by uniform mutation to the original reference. We note that *L*(*s*_*o*_, *s*_*m*_) may occasionally be sub-optimal, that is there may exist *s′*_*o*_ such that *L*(*s′*_*o*_, *s*_*m*_) *< L*(*s*_*o*_, *s*_*m*_) where *in silico* mutations introduced resulted in a sequence more similar to some other sequence in the database than the original.

For real datasets, 257 raw read sets that produced full-length contigs for at least one segment were chosen from the sequencing runs described above. The assembled contigs were annotated with BLAST (sorting by e-value, bit-score, and length), and raw reads classified by VAPOR. The percentage identity of VAPOR classifications to each contig was recorded.

### Detection of Reassortments and Zoonotic Strains

For assessment of reassortment classification, two simulations were performed. Firstly, 9659 avian, 18,308 human, and 2893 swine complete influenza genome sets were downloaded from the NIVR. 250 human genome sets were randomly selected. Another 250 were randomly selected with a single segment swapped with a randomly chosen avian or swine segment. For each, 1000 reads from each segment were simulated uniformly with an error rate of 0.5%. Each set of reads was classified with VAPOR. For the reference strains chosen by VAPOR for each segment, respective HA sequences were compared by global alignment, and percentage identity (PID) taken. If the maximum pairwise distance between chosen strain HA sequences exceeded a given threshold *v*, a classification of true was returned. Receiver operating characteristic (ROC) curves were generated by varying the parameter *v*. For assessment of intra-subtype reassortment classification, the same experiment was performed with randomly chosen H3N2 genomes.

### Computational Resources

In all cases, experiments were performed natively on a 96 core, 1.4 TB memory CentOS version 7.4.1708 virtual machine hosted by CLIMB [38], with GNU parallel [39] where required.

## Results

### Benchmarking Single-Reference Mapping

A range of mapping programs (Minimap2, BWA-MEM, Hisat2, and NGM) were compared to assess possible data loss when single references are chosen for mapping of short reads from influenza virus WGS datasets. For the first experiment, simulated reads from 16,679 human, 552 avian, and 4054 swine H1N1 HA sequences retrieved from the NIVR were mapped to the reference strain A/California/07/2009 (H1N1). Reads were simulated with an additional 0.05% error on top of simulated sequencing error to account for the combined effect of intra-host population variation and RT-PCR error. This error rate was found to be conservative when compared to the raw error rate in our datasets, as shown by Supplementary Figure 2, which was frequently higher than 2%. The proportion of successfully mapped reads for each tool and host species is given in Figure 2. In this case, using a single reference strain with any of the programs resulted in unmapped reads. NGM resulted in the lowest average percentage of unmapped reads. When utilizing a database of all H1N1 sequences from human hosts, Minimap2, NGM, BWA-MEM, and Hisat2 had mean mapping percentages of 87.2, 92.2, 89.1, and 84.9% respectively; as such, even for these influenza sequences, data loss was not uncommon, possibly due to samples in the database representing human infection from zoonotic strains. However, for avian and swine samples, read recovery was poor. For NGM, only 34.1% of avian reads mapped successfully on average. Swine sequences were mapped with intermediate success. This provides evidence that, should zoonotic strains be sequenced in routine surveillance, they may fail to map entirely, and go undetected. We note that this analysis is not an evaluation of overall mapping performance, since such an analysis must include mapping scores, but evidence that regardless of software, data loss may potentially occur.

**Figure 2:**
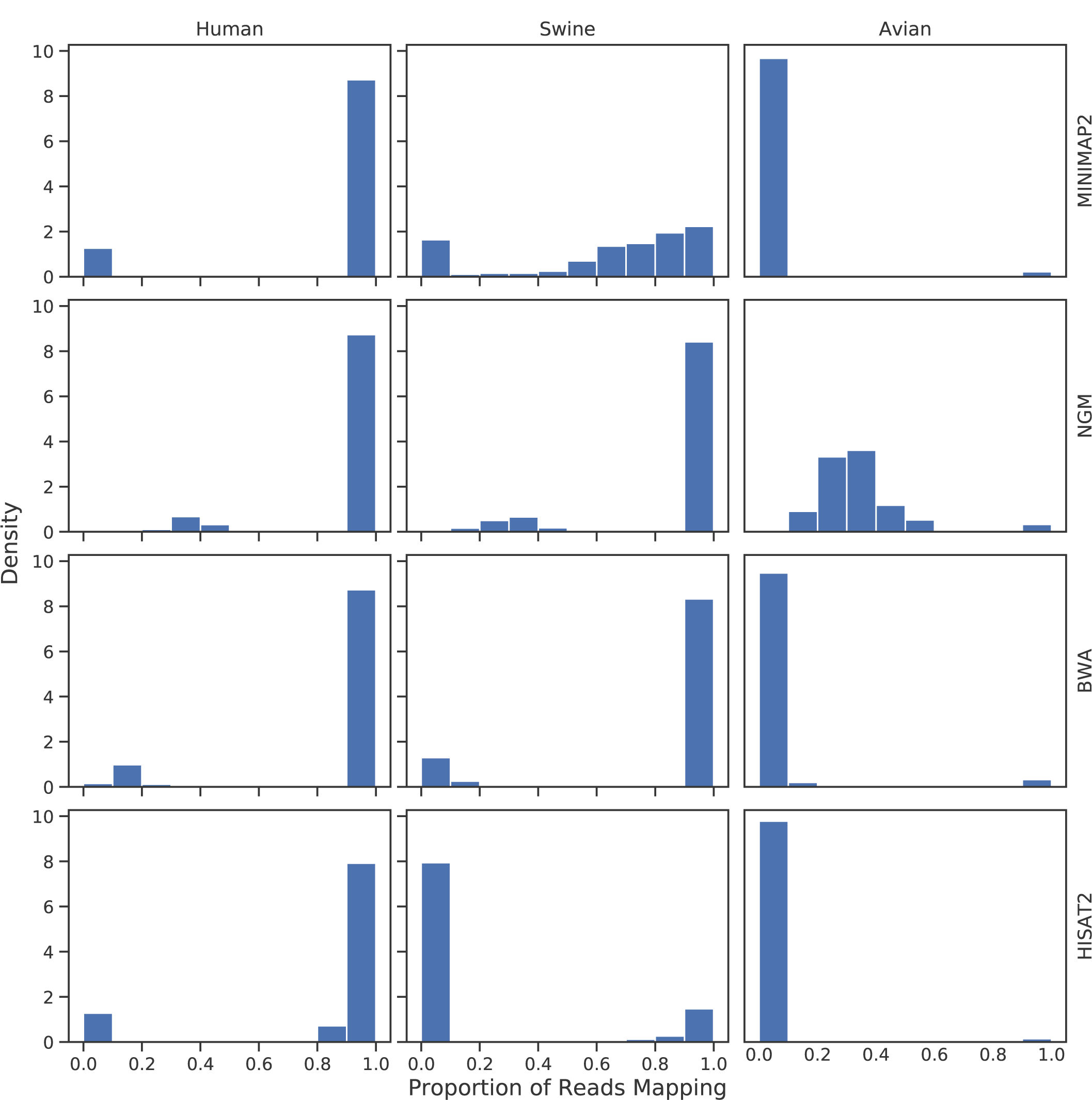
Density histograms showing proportion of mapped reads in samples, by software and dataset. Reads were simulated for each dataset retrieved from the NIVR: 16,679 Human H1N1 HA (left column); 552 avian H1N1 HA (middle column); 4054 Swine H1N1 HA (right column). All sequences were mapped to California/07/2009. For human sequences, most simulated datasets mapped successfully, although even for this dataset, around 10% of samples had some proportion of unmapped reads. However, for avian and swine sequences, mapping quality was poor, and often failed entirely. Even for the best performing software, NGM, avian sequences in particular mapped poorly.

Secondly, in order to assess how read recovery varies with sequence divergence, reads were simulated by taking the coding sequence of A/Perth/16/09 HA and subjecting it to *in silico* uniform mutation at specified rate, with additional read error of 0.05% as before. These results, shown in Figure 3, demonstrate that, for all mapping programs, at approximately 10% mutation, read recovery begins to regularly diminish, which is insufficient for robust mapping of influenza strains from different species. Furthermore, for several of the programs tested, mapping quality was suboptimal beyond 1-3% mutation.

**Figure 3:**
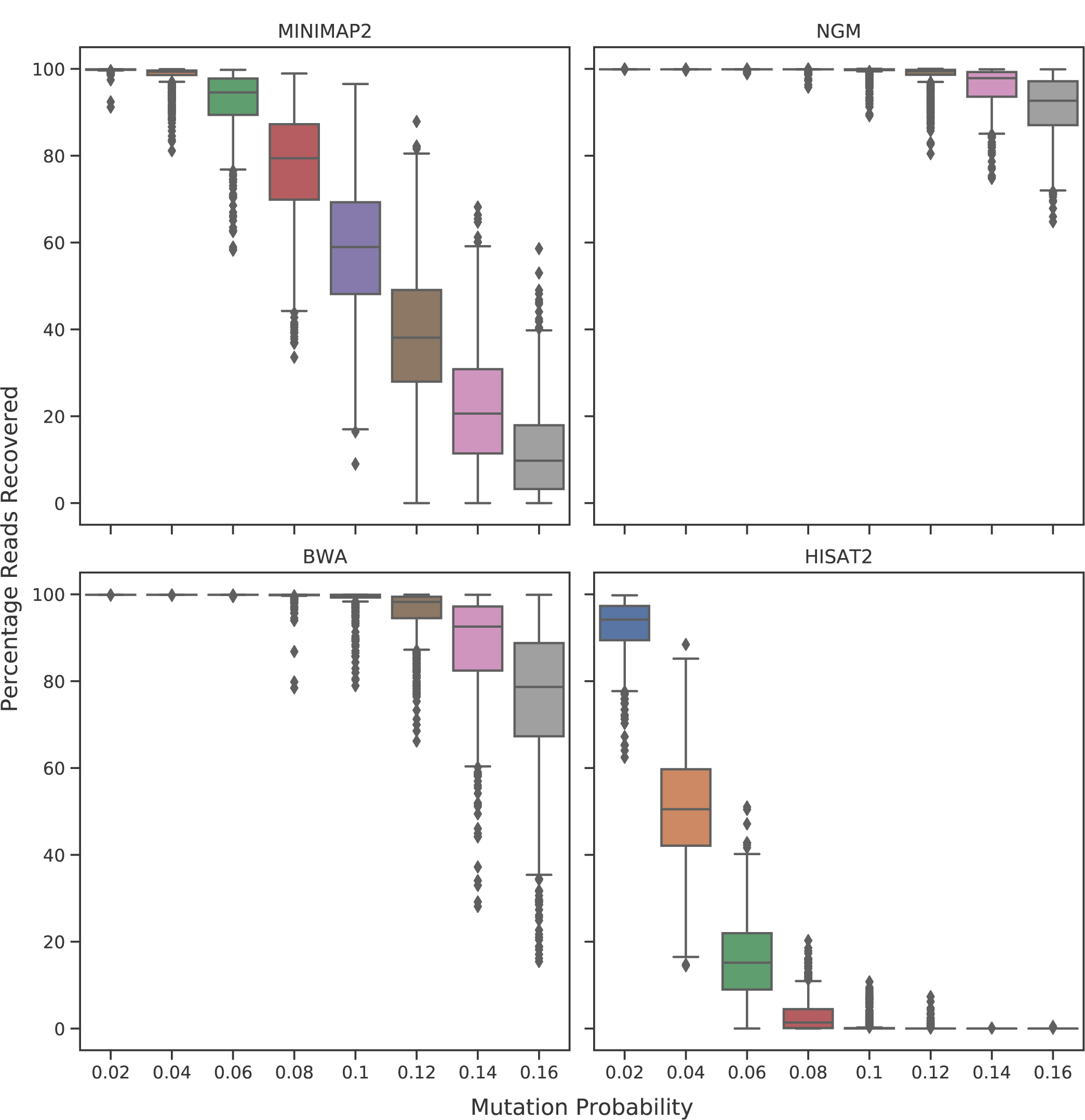
Box plots showing percentage of simulated Human H3N2 HA reads mapping to Perth/16/2009 for each software, with *in silico* uniform mutation at indicated per-base probability. Reads were simulated from *in silico* uniformly mutated Perth/16/2009 HA with the indicated per-base probability, approximately corresponding to 2 to 16% divergence. Reads were additionally subjected to 0.05% substitution to account for technical noise, such as from RT-PCR, and biological noise, such as from intrahost variation. Data loss was frequently observed with all tools beyond 10% mutation. Outliers are indicated as diamonds. N=1000 for each category.

### Classification Performance Simulation

In order to assess the performance of classification from simulated reads, our tool, VAPOR, was compared to MASH and consensus BLAST classification. Reads were simulated from randomly selected NIVR H1N1 HA sequences mutated with a given uniform per-base probability, with additional read error of 1% to provide challenging datasets. A fourth category included simple simulated intra-host populations (denoted as 3%/Q). Figure 4 shows the additional Levenshtein distance, *L*_*A*_ = *L*(*s*_*m*_, *s*_*c*_) *- L*(*s*_*m*_, *s*_*o*_), for each tool, where *s*_*o*_, *s*_*m*_, and *s*_*c*_ are the original, mutated, and retrieved database sequences respectively. Mean coverage for simulated reads was 77.76 for single-sequence simulations, and 96.03 for simulated intra-host populations. The average additional distance of retrieved sequences for MASH were 4.69, 5.24, 6.83, and 7.28, showing some sensitivity to additional simulated variant noise; for all cases mean additional distance for BLAST and VAPOR were below 0.74 and 0.88 respectively. For MASH, the 75%, 95%, and 99% percentiles for retrievals for the 3% threshold were 11.00, 24.00, and 37.04. However, for BLAST and VAPOR, these percentiles were under 12 and 14 respectively for all cases. These results show that references chosen by BLAST and VAPOR were often near-optimal or optimal, despite a large amount of noise, and that the performance difference between these approaches was very small. These results show that the algorithm used by VAPOR facilitates accurate classification of influenza strains directly from reads, comparable in accuracy but faster than BLAST for WGS read sets, which is generally not computationally tractable for datasets with millions of reads.

**Figure 4:**
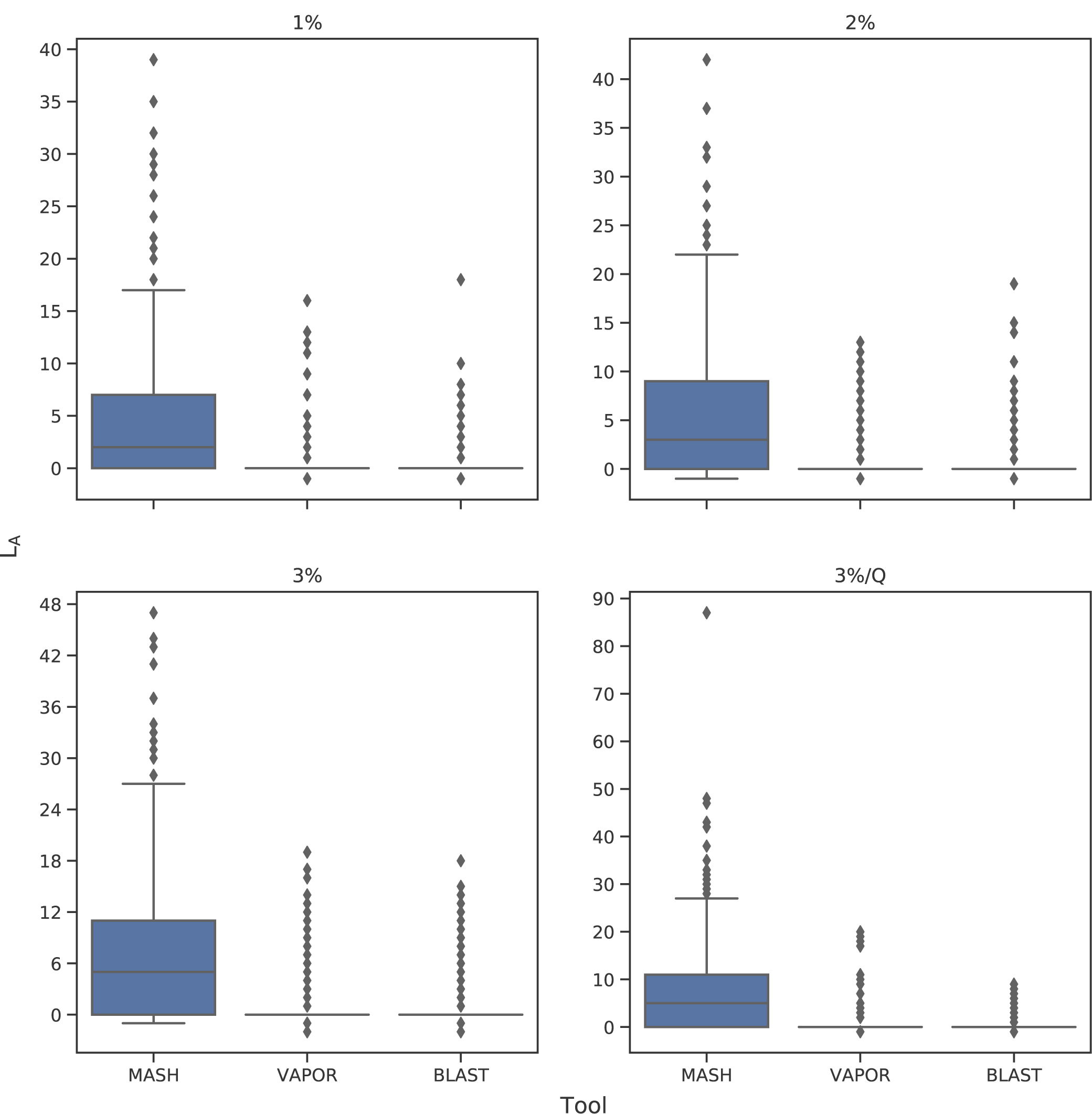
Box plots showing additional Levenshtein distance *L*_*A*_ = *L*(*s*_*m*_, *s*_*c*_) *- L*(*s*_*m*_, *s*_*o*_) of input sequence to output reference chosen by VAPOR, MASH, and BLAST consensus classification. Reads with 1% error rate were generated from randomly selected references mutated *in silico* by 1%, 2%, 3% and 3% with additional biological intra-host variant noise simulation 3%/Q, and repeated 500 times for each category. *L*_*A*_ is defined as Levenshtein distance of a classified sequence *s*_*c*_ to original mutated sequence *s*_*m*_, minus the distance of the original mutated sequence *s*_*m*_ to the original non-mutated reference sequence *s*_*o*_. Outliers are indicated as diamonds. Performance of VAPOR was generally equivalent to that of BLAST. For both of these tools, classification most often resulted in none, or a few extra incorrect bases. Sequences ranked highest by MASH were often sub-optimal.

### Real Data Classification Performance

Unlike BLAST and MASH, VAPOR can be applied directly on reads with no pre-processing. As such, in order to validate the performance of VAPOR directly on real datasets, we took raw reads from 257 samples corresponding to 1495 segment contigs previously processed and assembled with IVA, with a single full length contig each previously annotated by BLAST. In each case, corresponding reads were classified by VAPOR. The chosen reference was compared by global alignment to the assembled full length contigs. Figure 5 gives a scatter plot showing the PID of references retrieved by VAPOR to the assembled contig versus the PID of references selected by BLAST classifications of contigs. Comparison to BLAST classification of contigs was used to provide a baseline near-optimal classification. The mean percentage identity between contig and VAPOR classification was 99.82%. In the case of NS1, VAPOR outperformed BLAST annotation of assembled contigs, with a mean of 99.48 versus 98.74. On closer inspection, this was a result of the method used to sort BLAST results. These results show that in most cases tested, VAPOR was able to accurately identify a sample from reads with comparable performance to BLAST annotation of assembled contigs. We note that, for some contigs, neither BLAST nor VAPOR could achieve classifications with a PID greater than 97%. Manual examination of these samples showed large deletions, with at least one a likely misassembly (deletion including start codon, inclusion of 5’ UTR).

**Figure 5:**
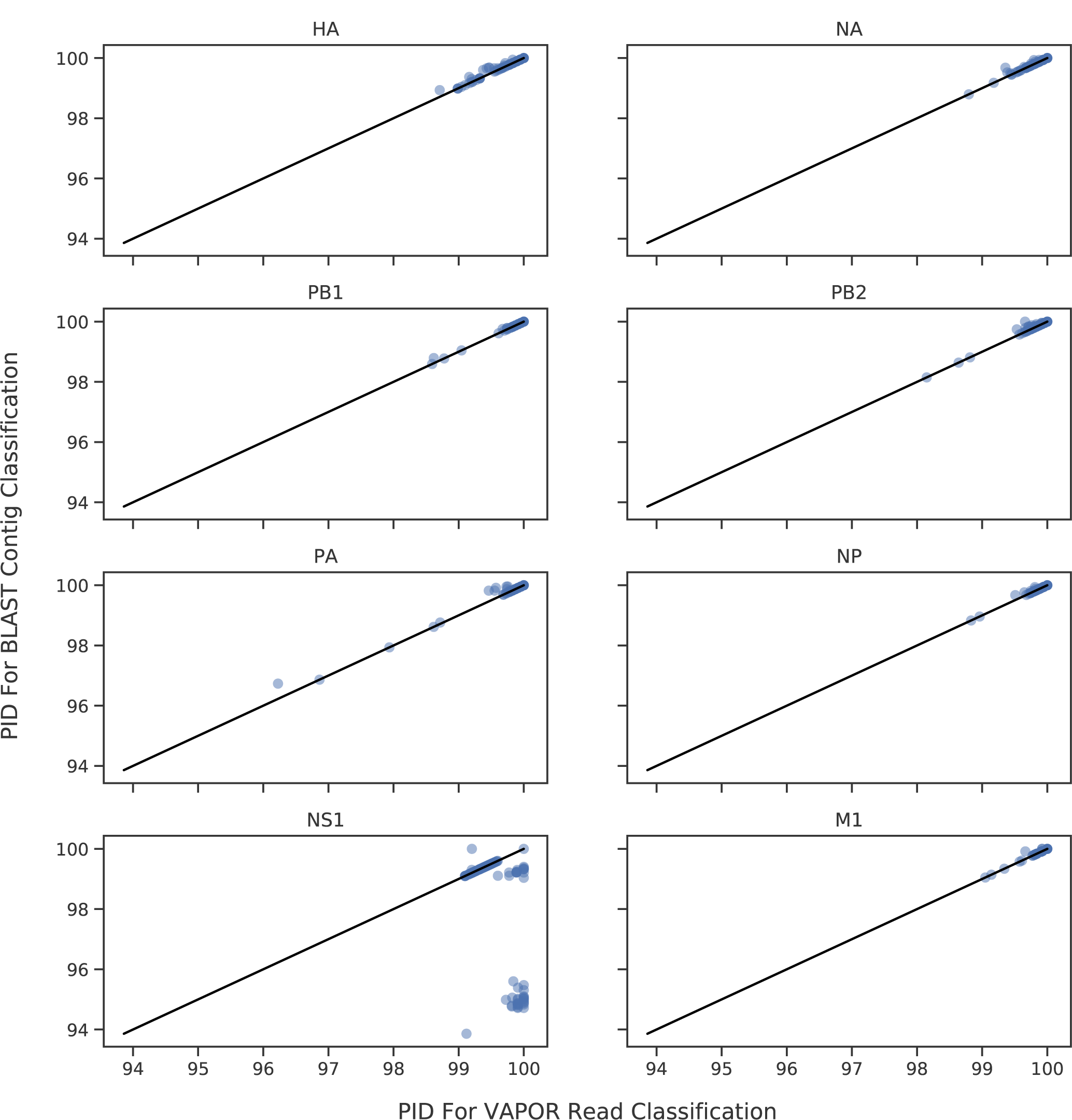
Scatterplots showing PIDs of VAPOR read classifications versus BLAST contig classifications with respect to assembled contigs for all 8 major segment coding sequences. Black lines indicate *x* = *y*. Points that fall below this line were classified better from reads with VAPOR. Points above the line were classified better with BLAST from contigs. VAPOR is capable in general of performing classification of reads to within 1% of the correct sequence. The mean PID of VAPOR classifications for all segments was 99.82%. For datapoints under 98% PID, BLAST was generally also not able of providing a better classification given the reference database.

### Mapping with Pre-Classification

In order to assess the utility of pre-classification for mapping, 25,533 full-length HA coding sequences from human, avian, and swine hosts were downloaded from the NIVR, and pairs were chosen randomly; one was used for read simulation, and the other as a mapping reference. In this case, mapping was performed both with and without pre-classification with VAPOR. Figure 6 gives the percentage of reads recovered against percentage identity for a pair, which shows that, for pairs chosen with less than 90% percentage identity, read recovery was poor. For mapping without pre-classification, mean recovery rates for Minimap2, NGM, BWA-MEM, and Hisat2 were 12.4%, 23.1%, 15.9%, 6.9%. However, with pre-classification using VAPOR, the mean was over 99.72% for all tools. These results demonstrate that mapping pipelines that include pre-classification are robust to sequences of zoonotic origin. Raw mapping performance was also assessed on real data by mapping datasets with Minimap2 with and without pre-classification. Figure 7 shows the number of additional reads mapped when pre-classification was performed. In all but one case, this resulted in a greater number of mapped reads, with a mean of 7816.03, corresponding to a mean percentage gain of 6.85%, including a case with over 68,000 additional reads. The maximum percentage increase was 13.32%. An outlier did occur where the number of mapped reads decreased. In this case, VAPOR identified several thousand more reads as influenza than were mapped. On further inspection, for this sample, reads mapped to both A/Perth/16/09 (H3N2) and A/California/07/09 (H1N1), indicating that the sample represented influenza from two different subtypes. As such, this sample may represent a true biological coinfection or a contamination, and could not be mapped to a single reference.

**Figure 6:**
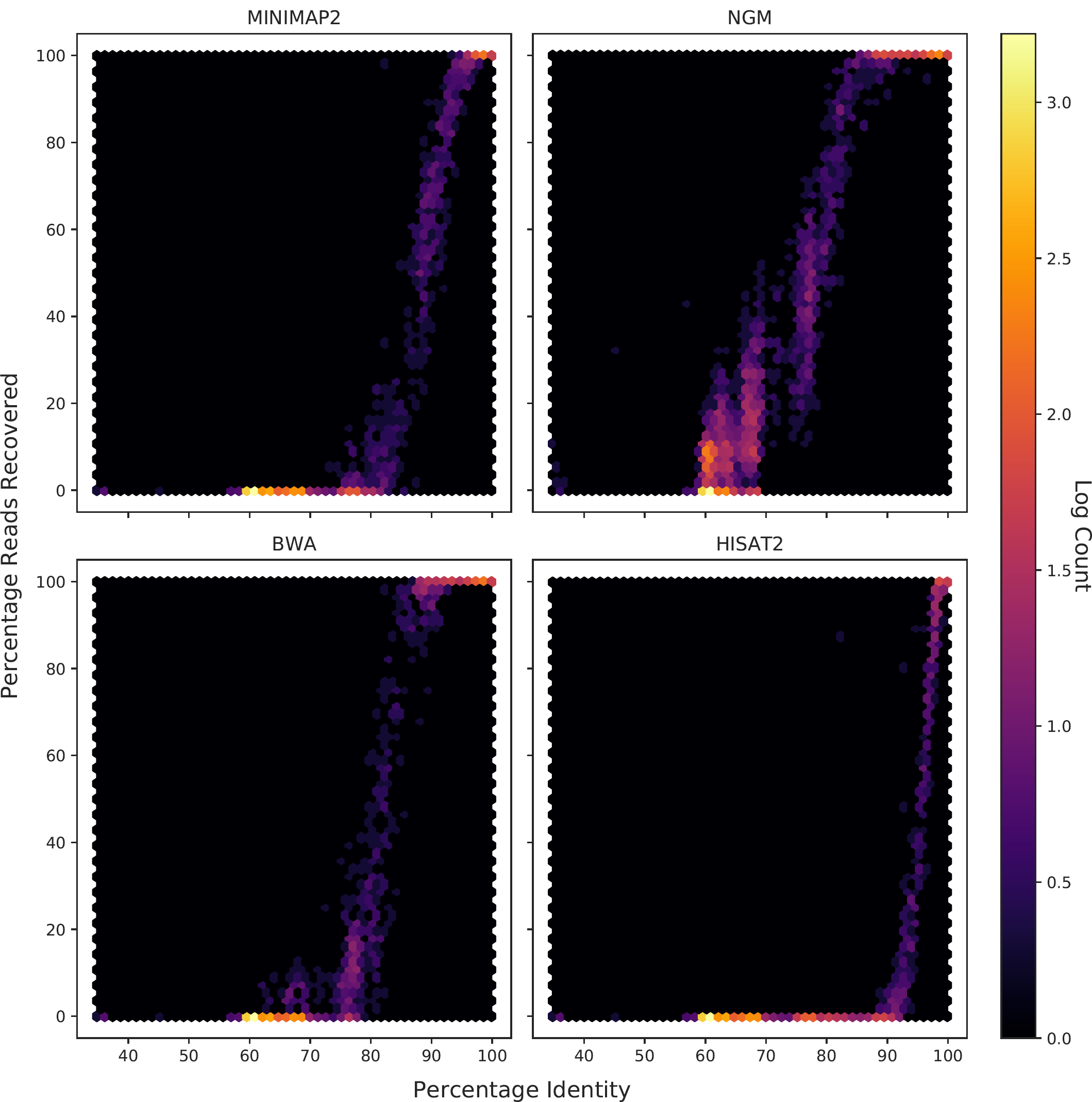
Read recovery versus percentage identity for randomly chosen influenza A HA sequence pairs. Influenza A HA sequences were chosen randomly from the NIVR database in pairs, one used as a reference, and one used to simulate reads for mapping. For percentage identities lower than 10%, mapping was unreliable. Furthermore, notably between strains of differing host origin, percentage identity was observed in many cases to be as low as 60%. These trends were observed for all tools, although performance dropped off more gradually for NGM.

**Figure 7:**
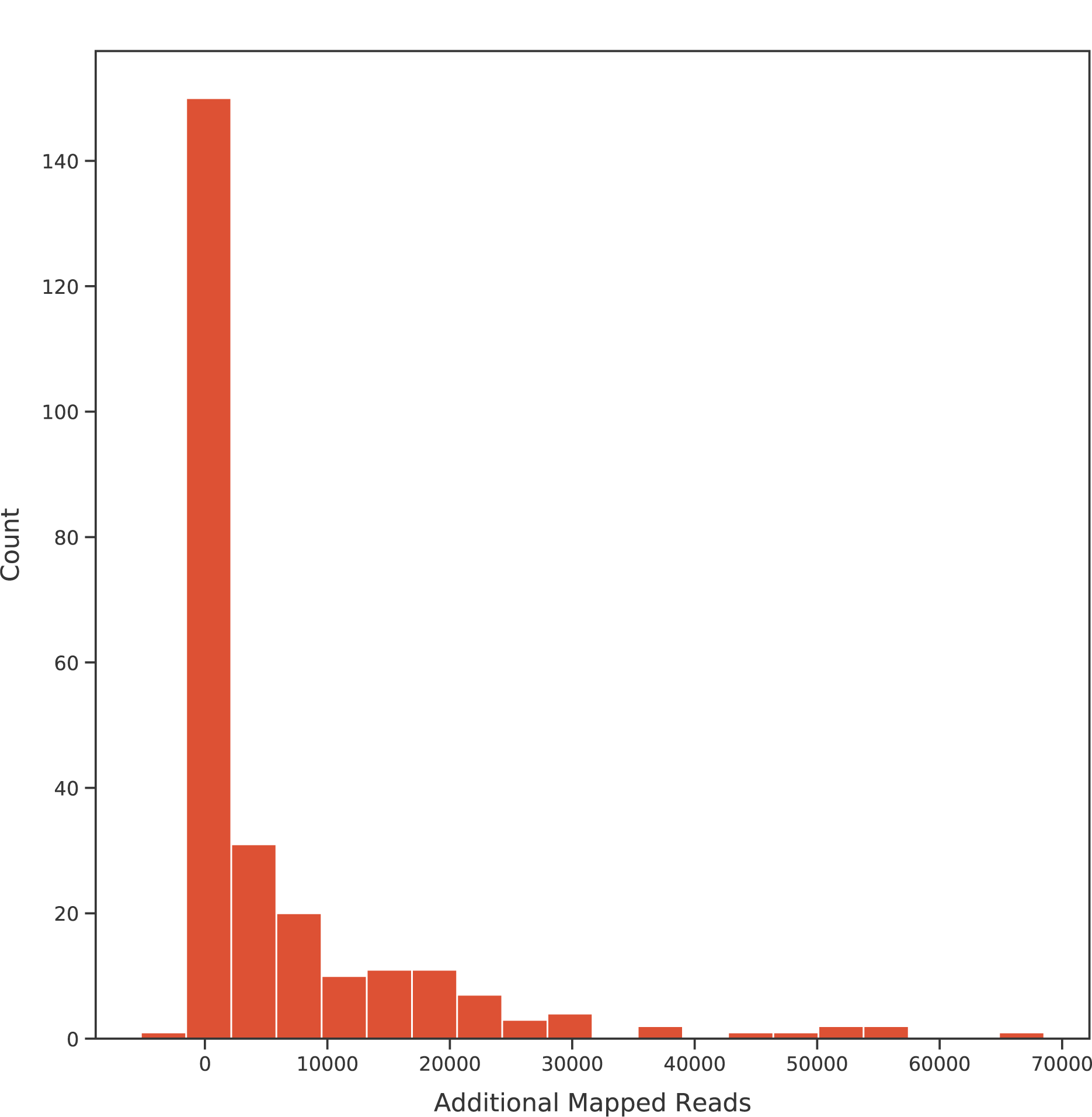
Additional Number of reads mapped by Minimap2 with VAPOR pre-classification for 257 real WGS datasets. Pre-classification with VAPOR on average resulted in 7816.03 more mapped reads. Several samples gained more than 50,000 reads by choosing a suitable reference. For one sample, representing a possible coinfection, 5221 fewer reads mapped when using a single reference chosen by VAPOR.

### Detection of Reassorted Strains Directly from Reads

In order to assess the application of read pre-classification to reassortment detection directly from reads, 250 simulated reassortment events with zoonotic strains were mixed with 250 complete genome sets, reads simulated, then classified by VAPOR. A simple reassortment classifier was used on the output of VAPOR, which compared the minimum pairwise PID of the HA sequences of the 8 strains assigned by VAPOR to each segment; if this PID was below a given parameter *v*, a reassortment was called. A ROC curve is shown in Figure 8, illustrating the performance of this classification strategy. Simulated zoonotic reassortments were detected with 97.2% true positive rate (TPR) and 0.08% false positive rate (FPR) for a *v* of 91.35%. This is expected because, as previously shown, VAPOR generally was able to classify strains to within a few base-pairs; randomly chosen zoonotic strains generally had PIDs of less than 90% to human strains, depending on origin. We note that, given the database used, some avian strains may have been isolated from humans, and labelled as human; as such, perfect classification with this dataset may be impossible. In order to provide a more difficult reassortment detection task, the same experiment was performed between human H3N2 sequences. We found at a PID threshold of 96.3%, a TPR of 76.8% could be achieved at a FPR of 10.8%. This result was expected given that sequences from different H3N2 strains generally have a PID within a few percent. In total, these results provide evidence that reassortments with zoonotic strains can be detected directly from reads with reasonable accuracy, but that intra-lineage reassortments may be more difficult.

**Figure 8:**
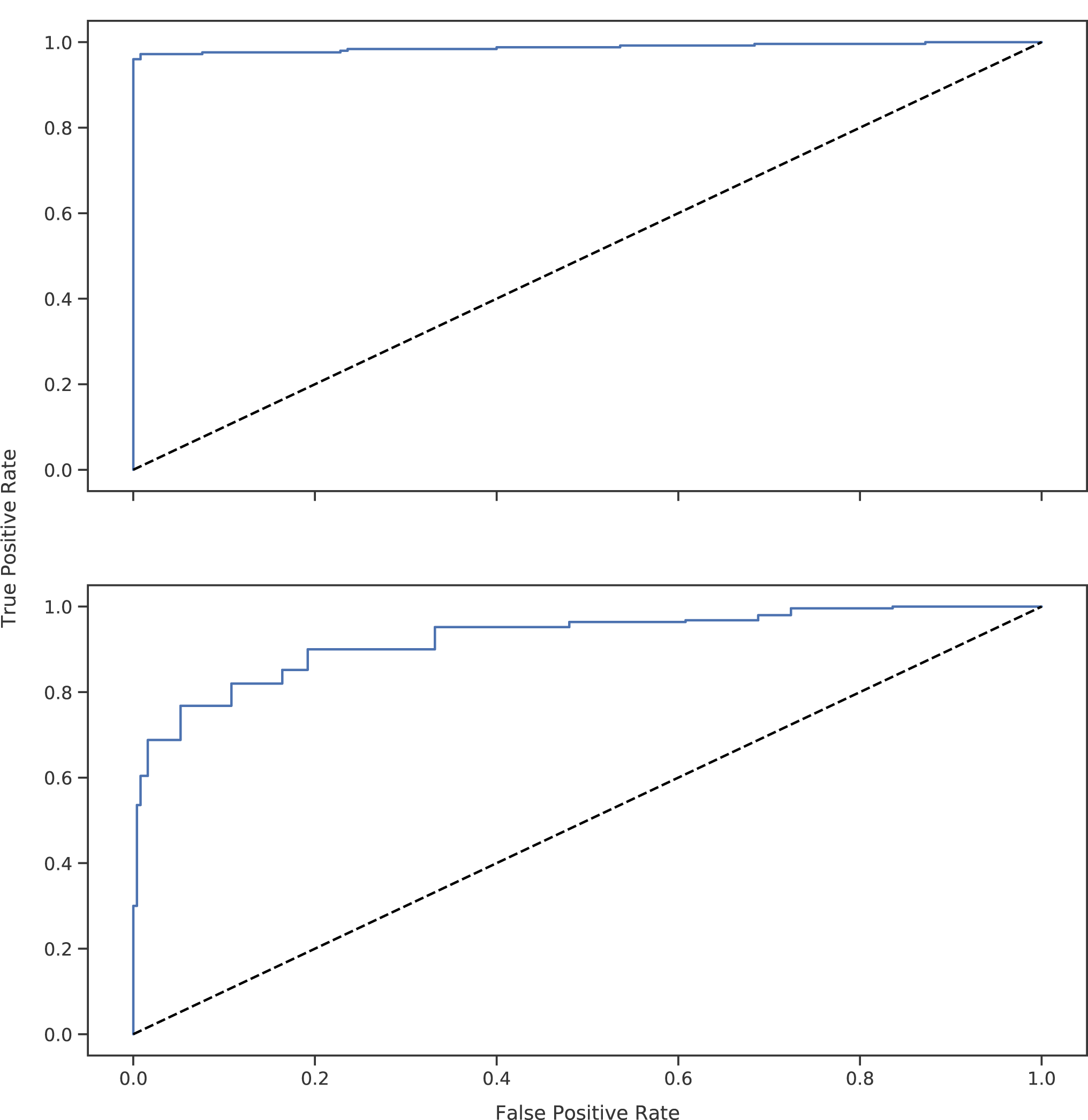
Receiver operating characteristic (ROC) curve for classification of simulated reas-sortment events. Although most zoonotic reassortments were detected at all parameter values (top), intra-H3N2 lineage (bottom) reassortments were more difficult to detect. The curve was generated by varying *v*, the minimum PID between VAPOR classifications of individual segment strains on the basis of HA. Due to the noise present in VAPOR classifications, as well as the close sequence similarity of H3N2 sequences, all parameter values with high TPR corresponded to a large FPR.

## Discussion

### Mapping Approaches and Improvement with VAPOR

We provide evidence that, in the best case, approaches for influenza virus analysis that use mapping to a single reference may result in data loss due to biological variation and noise. As shown in Supplementary Figure 1 and 2, influenza strains continually accumulate substitutions relative to a single reference (approximately 5 substitutions per year for H3N2), and reads may have a high error rate (>2%). Often, mapping to a single reference may be most unreliable for important samples, such as zoonotic transmission events. In the worst cases, mapping may fail completely, when usable data is present, requiring time and expertise to resolve with more complex methods. Our approach largely avoids these pitfalls altogether, allowing much simpler pipelines and alignment visualization via standard genome browsers, while also retaining the advantages of using a mapping-based approach for analysis. We chose Minimap2, BWA-MEM, NGM, and Hisat2 in order to represent a range of mapping softwares. BWA in particular has found use in general for influenza read mapping [9][17][22][40][41][42][43]. In other cases, software such as Bowtie2[44] have been used [10][16] for mapping to single references. In some cases, references were chosen by mapping-based approaches for selection [22]. Of these softwares, only NGM was developed with specific robustness to variation. Furthermore, the experiments reported were not intended as complete evaluations of the programs, since such an evaluation must also include mapping quality. Our data, however, does demonstrate that pre-classification with raw reads provides a broad strategy to improve robustness of pipelines and achieve faster results. For the chosen references, A/Perth/16/2009 (H3N2) and California/07/2009 (H1N1) were chosen as vaccine strains recommended by the WHO multiple times, and have also been used previously as references [9]. In other cases, different single references have been used [10]. They represent single strains that are well known, and may be used to represent each subtype. We do not believe that using different individual strains would affect the trends demonstrated.

We note that alternative approaches exist, including mapping to a large sequence database, but this does not allow for visualization of an alignment, and subsequent analysis such as characterization of point mutations. We note that in principle, pre-classification with any software could work reasonably well. MASH performed well in simulations. However; using an optimal reference is ideal, since for later advanced applications, such as transmission events, or study of intra-host variation, the closest possible reference may be necessary. Furthermore, VAPOR permits simultaneous filtering out of any non-human or bacterial reads with optimal reference selection. Whilst BLAST performed well for individual read classification, it is often too slow for general application. With regards to *de novo* assembly, in the cases where not enough initial data exists to assemble fragments, mapping allows analysis of limited fragments. Furthermore, assembly of virus genomes can be slow, often taking several days for a single sample when contaminant reads - such as human DNA are present. Finally, misassembly can occur [21].

In all but one of our real data cases examined, pre-classification with VAPOR resulted in a greater number of mapped reads than mapping to 4 reference strains from A/H3N2, A/H1N1, B/Victoria, and B/Yamagata. However, for a single sample, which contained influenza sequences from two clades, the number of mapped reads was reduced. Although VAPOR can report the number of influenza sequences detected in total, future study should be utilized to develop methods of coinfection detection. In these relatively rare cases, a single reference is not sufficient for mapping.

### VAPOR Algorithm and Performance

We have presented a novel approach to virus classification from short reads data using DBGs. In future study, as public sequence data accumulates, our algorithm may show promise in WGS approaches for other RNA viruses with small genomes, such as measles virus, Hepatitis C Virus, Human Immunodeficiency Virus (HIV), or Ebola virus. In general, our approach may have applications to short, variable genomes with high redundancy databases. We have shown that in many cases, VAPOR outperforms MASH, and has comparable performance to BLAST-based approaches. Furthermore, the algorithm used by VAPOR is well suited to simultaneous pre-filtering of contaminating human or bacterial sequences in samples, although we note that, in cases of coinfection, our algorithm may not be sufficient. Lastly, improved speed may be achieved by future implementation in C++, although generally, for the datasets examined, VAPOR can run within 5 minutes on a laptop with a 2.60GHz i7-6600U CPU.

Several default parameters were explored during development, but not exhaustively. A kmer size of 21 was utilized, as this was also able to perform read pre-filtering from contaminating sequences, without addition of a separate parameter. Similarly, parameters controlling the minimum fraction of required kmers for seed extension, as well as the top percentile of seeds chosen for extension could be adjusted, possibly to improve speed. However, in the read sets examined, the default parameters were generally sufficient to ensure matches were found, and did not appear to exclude potentially optimal matches. However, for novel strains that differ greatly from all strains previously observed, more sensitive parameterizations may be required.

### Real-data Classification

As shown in Figure 6, we note that the BLAST contig classification strategy we used performed poorly on NS1. This was due to sorting by e-value, bit-score, and length over percentage identity, combined with the presence of some NS1 sequences in the database which were longer than the required coding region. We opted to include this result to illustrate a potential pitfall that can occur with automated BLAST classification. Although sorting by PID may alleviate this problem, it may also yield shorter, incomplete alignments. For some samples, neither BLAST nor VAPOR could retrieve a sequence closer than 96% to the assembled contig. For some samples, this was due to large deletions present in the assembled contig. Although some of these deletions may be present in the true biological sequences, for at least one, this was due to suspected misassembly. These assemblies were also included to draw attention to potential problems that may be encountered during analysis. Furthermore, samples with deletions of ambiguous origin could not be excluded.

### Reassortment Classification

Over 97% of simulated zoonotic transmission events or reassortments could be identified at a cost of a 0.08% FPR using a simple alignment strategy whereby the PID of the HA sequences corresponding to the strains assigned to each segment are compared. Whilst some false positives occurred, this strategy provides a basis for pre-screening that can then be confirmed with slower methods as required. Intra-subtype classification from a single host species, such as human H3N2 was more difficult to classify. In this case, reported positives could be further validated by slower methods such as phylogenetic placement of assembled contigs. We note that it is not known *a priori* if any of the NIVR genomes are reassortments themselves. It is also possible that randomly choosing zoonotic strains to reassort is not biologically accurate, since there may be a limit on the similarity of reassorted sequences. However, we applied the same methodology to H3N2 sequences in order to demonstrate feasibility in detecting reassortment between very similar strains, although this was less accurate.

### Conclusions

Here we demonstrate that influenza sequence pre-classification with VAPOR allows alignment visualization, minimizes data loss, reduces pipeline complexity, and allows for classification of zoonotic strains and reassortments directly from reads. We believe that the simplicity of our approach has potential to alleviate several difficulties associated with current bioinformatics pipelines, and could reduce workloads in public health surveillance. Lastly, whilst we have tested VAPOR extensively for use with influenza, we believe our approach may be more broadly applicable to other sequence data, particularly small RNA and DNA viruses.

## Abbreviations

HA: hemagglutinin
NA: neuraminidase
WGS: whole genome sequencing
WHO: world health organization
RT-PCR: reverse-transcription polymerase chain reaction
NIVR: NCBI Influenza Virus Resource
GISAID: Global initiative on sharing all influenza data
NGS: next-generation sequencing
PHW: Public Health Wales
wDBG: weighted *de* Bruijn graph

## Availability and Requirements

1. Project name: VAPOR
2. Project home page: https://github.com/connor-lab/vapor
3. Operating system: Any with a command line and python3
4. Other requirements: NumPy >= 1.51.1
5. Programming language: python 3.x
6. License: GNU GPL 3.0
7. Restrictions to non-academic use: None

## Availability of Data and Materials

All scripts and pipelines used for simulations can be found https://github.com/connor-lab/ in the following repositories: `vapor benchmark mapping; vapor benchmark simulation; vapor benchmark realdata; vapor benchmark simulation`. Short read data can be found at https://s3.climb.ac.uk/vapor-benchmark-data/vapor_benchmarking_realdata_reads_filtered_18_03_18.tar. Human sequences were depleted from this data as described in Methodology. All other data required for reproduction of results can be obtained according to the instructions found in the respective repositories.

## Acknowledgments

We would like to acknowledge and offer our thanks to the Pathogen Genomics Unit (PenGU) at Public Health Wales for their efforts and assistance in performing laboratory work, without which this research could not have been performed. Likewise, we would also like to thank Angela Marchbank and the Sequencing Hub at Cardiff University for the sequencing work performed.

## References

[1] Taubenberger JK, Kash JC. Influenza virus evolution, host adaptation, and pandemic formation. Cell Host and Microbe. 2010;7:440–451.

[2] Petrova VN, Russell CA. The evolution of seasonal influenza viruses. Nature Reviews Microbiology. 2018;47:47–60.

[3] Tafalla M, Buijssen M, Geets R, Vonk Noordegraaf-Schouten M. A comprehensive review of the epidemiology and disease burden of influenza B in 9 European countries. Human vaccines and immunotherapeutics. 2016;12:993–1002.

[4] Iuliano AD, Roguski KM, Chang HH, Muscatello DJ, Palekar R, Tempia S, et al. Estimates of global seasonal influenza-associated respiratory mortality: a modelling study. The Lancet. 2018;391:1285–1300.

[5] Sautto GA, Kirchenbaum GA, Ross TM. Towards a universal influenza vaccine: different approaches for one goal. Virology journal. 2018;15:17.

[6] Bouvier NM, Palese P. The Biology of influenza Viruses. Vaccine. 2008;26:D49–D53.

[7] Holmes EC, Ghedin E, Miller N, Taylor J, Bao Y, St George K, et al. Whole-Genome Analysis of Human influenza A Virus Reveals Multiple Persistent Lineages and Reassortment among Recent H3N2 Viruses. PLoS Biology. 2005;3:e300.

[8] McGinnis J, Laplante J, Shudt M, George KS. Next generation sequencing for whole genome analysis and surveillance of influenza A viruses. Journal of Clinical Virology. 2016;79:44–50.

[9] Rutvisuttinunt W, Chinnawirotpisan P, Simasathien S, Shrestha SK, Yoon IK, Klungthong C, et al. Simultaneous and complete genome sequencing of influenza A and B with high coverage by Illumina MiSeq Platform. Journal of Virological Methods. 2013;193:394–404.

[10] Meinel DM, Heinzinger S, Eberle U, Ackermann N, Schönberger K, Sing A. Whole genome sequencing identifies influenza A H3N2 transmission and offers superior resolution to classical typing methods. Infection. 2018;46:69–76.

[11] Zhou B, Lin X, Wang W, Halpin RA, Bera J, Stockwell TB, et al. Universal influenza B Virus Genomic Amplification Facilitates Sequencing, Diagnostics, and Reverse Genetics. Journal of Clinical Microbiology. 2014;52:1330–1337.

[12] Zhou B, Donnelly ME, Scholes DT, St George K, Hatta M, Kawaoka Y, et al. Single-Reaction Genomic Amplification Accelerates Sequencing and Vaccine Production for Classical and Swine Origin Human influenza A Viruses. Journal of Virology. 2009;83:10309–13.

[13] Houlihan CF, Frampton D, Ferns RB, Raffle J, Grant P, Reidy M, et al. Use of Whole-Genome Sequencing in the Investigation of a Nosocomial influenza Virus Outbreak. Journal of Infectious Diseases. 2018;218:1485–1489.

[14] Bao Y, Bolotov P, Dernovoy D, Kiryutin B, Zaslavsky L, Tatusova T, et al. The influenza virus resource at the National Center for Biotechnology Information. Journal of Virology. 2008;82:596–601.

[15] Shu Y, McCauley J. GISAID: Global initiative on sharing all influenza data - from vision to reality. EuroSurveillance. 2017;22:30494.

[16] Goldstein EJ, Harvey WT, Wilkie GS, Shepherd SJ, MacLean AR, Murcia PR, et al. Integrating patient and whole-genome sequencing data to provide insights into the epidemiology of seasonal influenza A(H3N2) viruses. Microbial Genomics. 2017;2018:4.

[17] Borges V, Pinheiro M, Pechirra P, Guiomar R, Gomes JP. INSaFLU: an automated open web-based bioinformatics suite from-reads for influenza whole-genome-sequencing-based surveillance. Genome Medicine. 2018;10:46.

[18] Wan Y, Renner DW, Albert I, Szpara ML. VirAmp: a galaxy-based viral genome assembly pipeline. GigaScience. 2015;4:19.

[19] Hunt M, Gall A, Ong SH, Brener J, Ferns B, Goulder P, et al. IVA: accurate *de novo* assembly of RNA virus genomes. Bioinformatics. 2015;31:2374–6.

[20] Orton RJ, Wright CF, Morelli MJ, King DJ, Paton DJ, King DP, et al. Distinguishing low frequency mutations from RT-PCR and sequence errors in viral deep sequencing data. BMC Genomics. 2015;16:299.

[21] Wymant C, Blanquart F, Golubchik T, Gall A, Bakker M, Bezemer D, et al. Easy and accurate reconstruction of whole HIV genomes from short-read sequence data with shiver. Virus Evolution. 2018;4:vey007.

[22] Yu X, Jin T, Cui Y, Pu X, Li J, Xu J, et al. Influenza H7N9 and H9N2 Viruses: Coexistence in Poultry Linked to Human H7N9 Infection and Genome Characteristics. Virology. 2014;88:3423–3431.

[23] Limasset A, Cazaux B, Rivals E, Peterlongo P. Read mapping on *de* Bruijn graphs. Bioinformatics. 2016;17:237.

[24] Holley G, Peterlongo P. Blastgraph: Intensive approximate patternmatching in sequence graphs and de Bruijn graphs. In: Stringology. 2012; p. 53–63.

[25] Liu B, Guo H, Brudno M, Wang Y. deBGA: read alignment with *de* Bruijn graph-based seed and extension. Bioinformatics. 2016;32:32243232.

[26] Salmela L, Rivals E. LoRDEC: accurate and efficient long read error correction. Bioinformatics. 2014;30:3506–3514.

[27] Altschul SF, Gish W, Miller W, Myers EW, Lipman DJ. Basic local alignment search tool. Journal of Molecular Biology. 1990;215:403–410.

[28] Ondov BD, Treangen TJ, Melsted P, Mallonee AB, Bergman NH, Koren S, et al. MASH: fast genome and metagenome distance estimation using MinHash. Genome Biology. 2016;17:132.

[29] Wood DE, Salzberg SL. Kraken: ultrafast metagenomic sequence classification using exact alignments. Genome Biology. 2014;15:R46.

[30] Li H. Minimap2: pairwise alignment for nucleotide sequences. Bioinformatics. 2018;34:3094–3100.

[31] Li W, Godzik A. Cd-hit: a fast program for clustering and comparing large sets of protein or nucleotide sequences. Bioinformatics. 2006;22:1658–1659.

[32] Li H, Durbin R. Fast and accurate short read alignment with Burrows-Wheeler transform. Bioinformatics. 2009;25:1754–60.

[33] Sedlazeck FJ, Rescheneder P, von Haeseler A. NextGenMap: fast and accurate read mapping in highly polymorphic genomes. Bioinformatics. 2013;29:2790–2791.

[34] Kim D, Langmead B, Salzberg SL. HISAT: a fast spliced aligner with low memory requirements. Nature Methods. 2015;12:357–6.

[35] Frampton M, Houlston R. Generation of Artificial FASTQ Files to Evaluate the Performance of Next-Generation Sequencing Pipelines. PLoS ONE. 2012;7:e49110.

[36] Li H, Handsaker B, Wysoker A, Fennell T, Ruan J, Homer N, et al. The Sequence Alignment/Map format and SAMtools. Bioinformatics. 2009;25:2078–9.

[37] Cock PJ, Antao T, Chang JT, Chapman BA, Cox CJ, Dalke A, et al. Biopython: freely available Python tools for computational molecular biology and bioinformatics. Bioinformatics. 2009;25:1422–1423.

[38] Connor TR, Loman NJ, Thompson S, Smith A, Southgate J, Poplawski R, et al. CLIMB (the Cloud Infrastructure for Microbial Bioinformatics): an online resource for the medical microbiology community. Microbial Genomics. 2016;2:e000086.

[39] Tange O. GNU Parallel - The Command-Line Power Tool.; login: The USENIX Magazine. 2011;36:42–47. Available from: http://www.gnu.org/s/parallel.

[40] Wu NC, Young AP, Al-Mawsawi LQ, Olson CA, Feng J, Qi H, et al. High-throughput profiling of influenza A virus hemagglutinin gene at single-nucleotide resolution. Scientific Reports. 2014;4:4942.

[41] Imai K, Tamura K, Tanigaki T, Takizawa M, Nakayama E, Taniguchi T, et al. Whole Genome Sequencing of influenza A and B Viruses With the MinION Sequencer in the Clinical Setting: A Pilot Study. Front Microbiol. 2018;9:2748.

[42] Leonard AS, McClain MT, Smith GJD, Wentworth DE, Halpin RA, Lin X, et al. Deep Sequencing of Influenza A Virus from a Human Challenge Study Reveals a Selective Bottleneck and Only Limited Intrahost Genetic Diversification. Virology. 2016;90:11247–11258.

[43] Jonges M, Welkers MRA, Jeeninga RE, Meijer A, Schneeberger P, Fouchier RAM, et al. Emergence of the Virulence-Associated PB2 E627K Substitution in a Fatal Human Case of Highly Pathogenic Avian influenza Virus A(H7N7) Infection as Determined by Illumina Ultra-Deep Sequencing. Virology. 2014;88:1694–1702.

[44] Langmead B, Salzberg SL. Fast gapped-read alignment with Bowtie 2. Nature methods. 2012;9:357–359.

