## Supplementary Figure 1 for "Influenza Classification from Short Reads with VAPOR Facilitates Robust Mapping Pipelines and Zoonotic Strain Detection for Routine Surveillance Applications"

### Divergence Over Time

In order to assess the accumulation of substitutions over time for purposes of mapping, 21007 H3N2 HA sequences were downloaded from the NCBI Influenza Virus Resource [1], and aligned with MAFFT [2]. As shown in Supplementary Figure 1, the hamming distance of aligned sequences from the earliest sequenced strain, A/Aichi/2/1968, was recorded. Outliers with more than 400 substitutions were excluded.

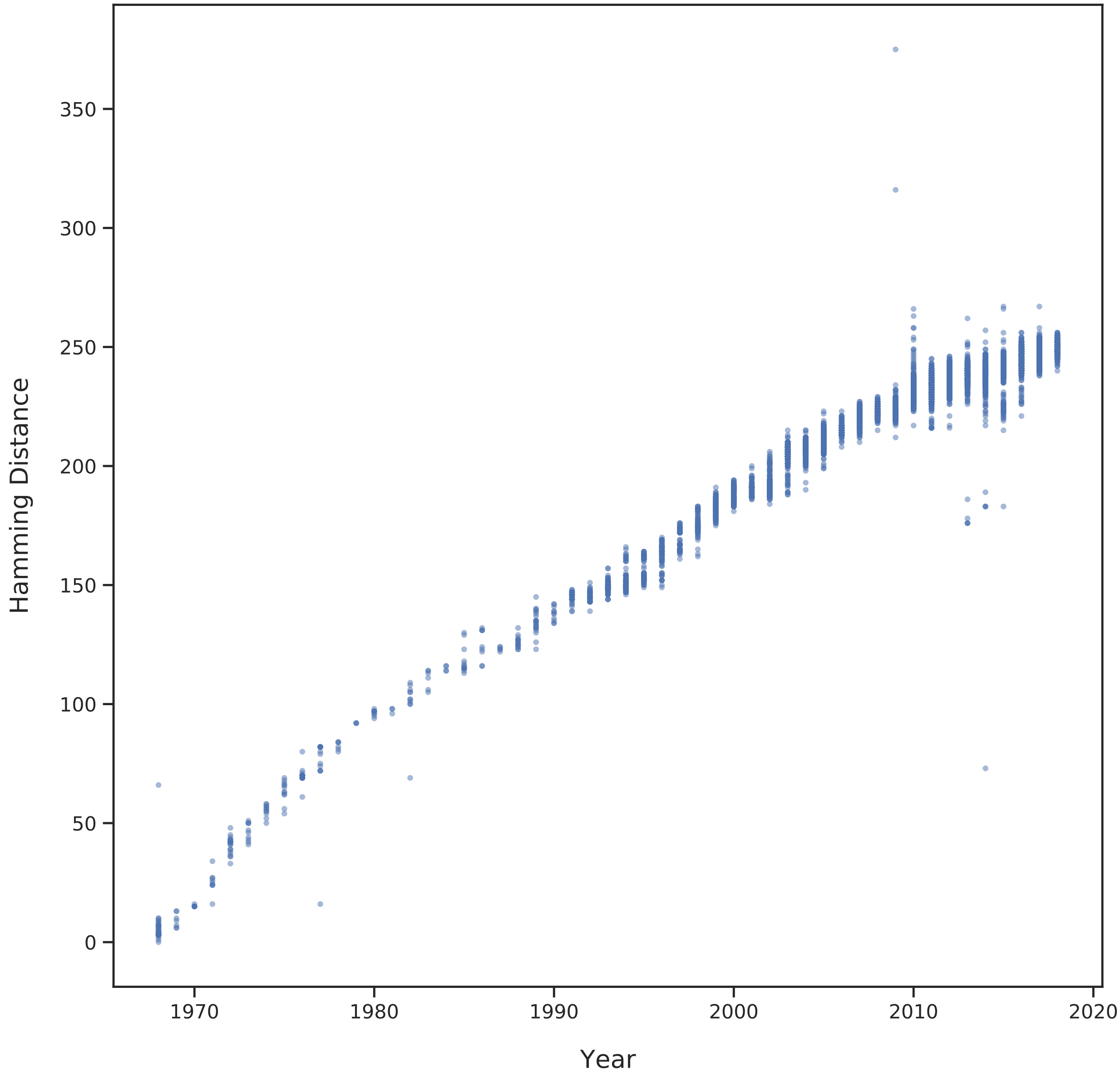


Figure 1: **Hamming distance of aligned H3N2 HA sequences from A/Aichi/2/1968**. On average, sequences accumulate a hamming distance of approximately 5 substitutions per year.
.

### References

[1] Bao, Y., Bolotov, P., Dernovoy, D., Kiryutin, B., Zaslavsky, L., Tatusova, T., Ostell, J., Lipman, D.: The influenza virus resource at the National Center for Biotechnology Information. Journal of Virology **82**, 596–601 (2008)

[2] Katoh, K., Standley, D.: MAFFT multiple sequence alignment software version 7: Improvements in performance and usability. Molecular Biology and Evolution **30**, 772–780 (2013)
