## Supplementary Figure 2 for "Influenza Classification from Short Reads with VAPOR Facilitates Robust Mapping Pipelines and Zoonotic Strain Detection for Routine Surveillance Applications"

### Read Error Distribution

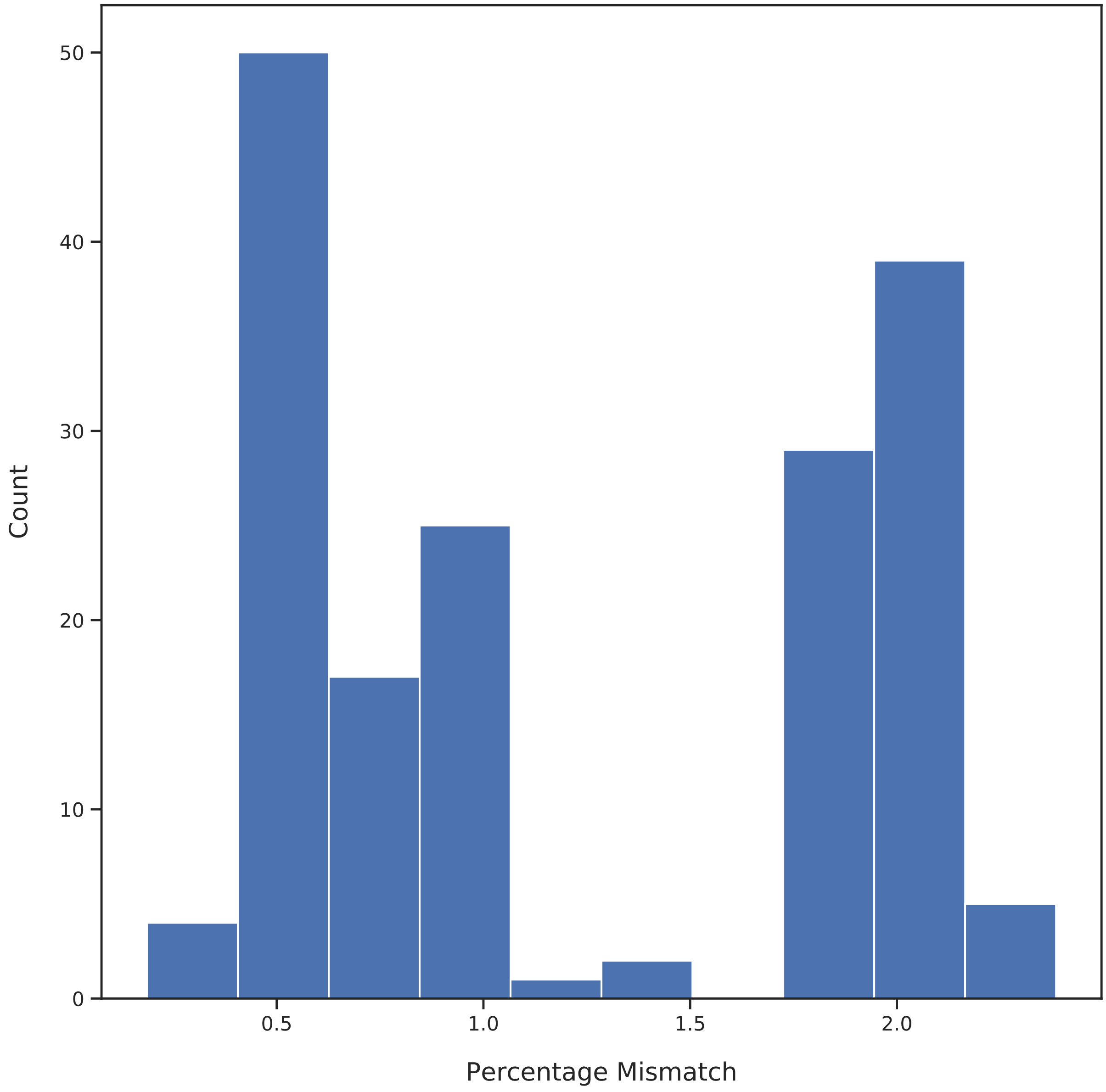


Figure 1: **Histogram giving percentage of mismatches in reads relative to assembled contigs**.
.
